# METASPACE: A community-populated knowledge base of spatial metabolomes in health and disease

**DOI:** 10.1101/539478

**Authors:** Theodore Alexandrov, Katja Ovchinnikova, Andrew Palmer, Vitaly Kovalev, Artem Tarasov, Lachlan Stuart, Renat Nigmetzianov, Dominik Fay, Key METASPACE contributors, Mathieu Gaudin, Cristina Gonzalez Lopez, Marina Vetter, John Swales, Mark Bokhart, Mario Kompauer, James McKenzie, Luca Rappez, Dusan Velickovic, Regis Lavigne, Guanshi Zhang, Dinaiz Thinagaran, Elisa Ruhland, Marta Sans, Sergio Triana, Denis Abu Sammour, Sarah Aboulmagd, Charlotte Bagger, Nicole Strittmatter, Angelos Rigopoulos, Erin Gemperline, Asta Maria Joensen, Benedikt Geier, Christine Quiason, Eric Weaver, Mridula Prasad, Benjamin Balluff, Konstantin Nagornov, Lingjun Li, Michael Linscheid, Carsten Hopf, Dimitri Heintz, Manuel Liebeke, Bernhard Spengler, Berin Boughton, Christian Janfelt, Kumar Sharma, Charles Pineau, Christopher Anderton, Shane Ellis, Michael Becker, József Pánczél, Georges Da Violante, David Muddiman, Richard Goodwin, Livia Eberlin, Zoltan Takats, Sheerin Shahidi-Latham

## Abstract

Metabolites, lipids, and other small molecules are key constituents of tissues supporting cellular programs in health and disease. Here, we present METASPACE, a community-populated knowledge base of spatial metabolomes from imaging mass spectrometry data. METASPACE is enabled by a high-performance engine for metabolite annotation in a confidence-controlled way that makes results comparable between experiments and laboratories. By sharing their results publicly, engine users continuously populate a knowledge base of annotated spatial metabolomes in tissues currently including over 3000 datasets from human cancer cohorts, whole-body sections of animal models, and various organs. The spatial metabolomes can be visualized, explored and shared using a web app as well as accessed programmatically for large-scale analysis. By using novel computational methods inspired by natural language processing, we illustrate that METASPACE provides molecular coverage beyond the capacity of any individual laboratory and opens avenues towards comprehensive metabolite atlases on the levels of tissues and organs.

## Introduction

Metabolites, lipids, and other small molecules play essential structural, energetic, and regulating roles in virtually all cellular processes ^1^. Their localization is specific to single cells, cell types, tissues, and organs, is regulated in time and space and at the same time regulates homeostasis in health and reprogramming in disease ^2^. Studying roles of metabolites in their native spatial context is a frontier of metabolism research that recently led to exciting discoveries in cancer ^3^, immunity ^4^, microbiome ^5^, and stem cells ^6^ research. However, spatial investigations of metabolism are challenging and can hardly be performed by using conventional metabolomics or imaging methods. Fueled by this demand, spatial metabolomics has emerged as an interdisciplinary approach combining untargeted detection of metabolites *in situ* with computational approaches aiding interpretation of the generated big spatial data.

A prominent technology for spatial metabolomics is imaging mass spectrometry (imaging MS) which samples mass spectra from a raster of pixels across a tissue section, with each mass spectrum representing relative molecular abundances at the corresponding pixel. Recent advances in surface sampling, mass spectrometry, and bioinformatics made imaging MS a technique of choice for spatial metabolomics in a variety of biomedical and pharmacological applications ^7–10^. Imaging MS has a near-single-cell spatial resolution of 1.4 μm on a commercial instrumentation ^11^, the throughput to analyze a tissue section in minutes ^12^, high specificity when using high-mass-resolution ^13^, sensitivity to detect small molecules at physiological micromolar concentrations^14^, and capacity to detect various classes of molecules including amino acids, central carbon metabolism intermediates, free fatty acids, various lipids, and neurotransmitters ^15^. Until recently, the progress and applications of imaging MS were hampered by the lack of algorithms for metabolite identification, namely for associating ion peaks and images with their corresponding molecules. We have recently addressed this issue and published a method for confidence-controlled metabolite annotation ^13^.

Here, we present METASPACE, a public knowledge base of spatial metabolomes in health and disease populated by a large community of users (Figure 1a). The cornerstone of METASPACE is a computational engine for metabolite annotation which searches for metabolites, lipids, and other small molecules in an imaging MS dataset. The engine estimates the False Discovery Rate (FDR) of metabolite annotations that provides quality control and – as demonstrated in other -omics ^16^ – makes annotated spatial metabolomes comparable between datasets, experiments, and laboratories ^13^. We created a user-friendly web app for data submission and for interactive exploration of annotated metabolite images. By sharing their results publicly, METASPACE users cooperatively created and continuously populate a knowledge base of spatial metabolomes.

**Figure 1.**
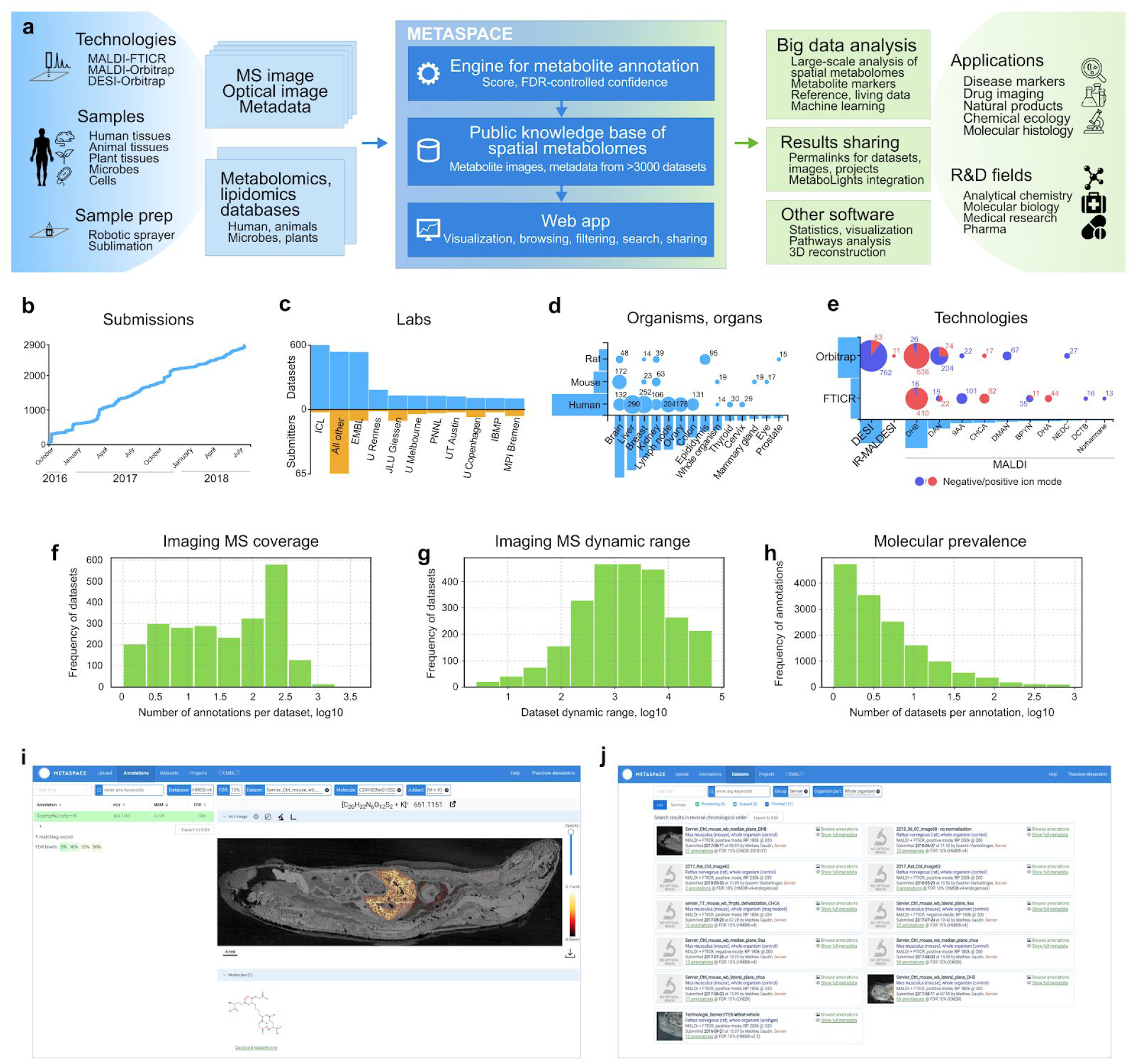
METASPACE is a community-populated knowledge base of spatial metabolomes from imaging MS data that integrates a cloud engine for metabolite annotation and a web app for visualization, exploration, and sharing annotated metabolite images. **A**. METASPACE concept and workflow. **B-E**. Number of submissions and properties of the submitted datasets. **F**. Histogram of the estimated coverage of imaging MS (number of annotations in a dataset at FDR 10%). **G**. Histogram of the estimated dynamic range of imaging MS (ratio between intensities of the most and least intense annotations, per dataset). **H.** Histogram of the prevalence of annotations (number of datasets where a molecule was annotated with FDR 10%). **I**. Screenshot of the METASPACE web app showing a distribution of glutathione in a mouse whole-body section contributed by Mathieu Gaudin, Servier (<u>link for the dataset, link for theshown metabolite image</u>). **J.** Screenshot of the web app showing all datasets from whole-body sections contributed by one laboratory, here the pharma company Servier (<u>link</u>).

## Results

### Engine

At the heart of METASPACE is a high-performance engine implementing our recent method for MS1-based metabolite annotation for imaging MS ^13^ (Figure 1a). The engine is open-source and exploits big data computing technologies for scalable parallel processing in cloud environment (https://github.com/metaspace2020). After receiving an imaging MS dataset in the imzML centroided format ^17^, the engine screens for molecules from over ten public metabolomics or lipidomics databases including HDMB ^18^, ChEBI ^19^, LIPID MAPS ^20^, SwissLipids ^21^ and provides molecular annotations on the level of molecular formula (Level 2 according to the Metabolomics Standards Initiative ^22^). Annotating a dataset of 10 GB takes below 10 minutes on a default cloud cluster. For each metabolite annotation, the engine provides its molecular formula, ion adduct, and metabolite ion image. Importantly, it also provides all isomeric molecular structures corresponding to this formula in the screened-for database thus helping understand the isomeric ambiguity of the results. The estimated Metabolite-Signal Match (MSM) score ranks the molecular formulas by their likelihood ^13^. The estimated FDR value quantifies the confidence of the results and makes them comparable between datasets. Importantly, FDR-controlled annotation serves as a quality control, as the engine provides no or only a few annotations for datasets of poor quality, e.g. when the mass spectra are misaligned or not properly calibrated ^13^. Overall, the engine enables rapid and comprehensive search for metabolites, lipids, and small molecules in imaging MS data and, through providing a yet missing capacity, bridges the gap between the imaging MS technology and biomedical applications.

### Knowledge base

The METASPACE knowledge base is a community-populated resource containing public results from the annotation engine, with every dataset including metabolite annotations, metabolite images, and associated metadata. Users can make the results public either during submission or later. As of January 2019, the knowledge base encompasses over 6 million metabolite images from over 140 contributors from over 70 laboratories who already submitted over 3200 public datasets from tissue sections from various organisms (human, mouse, rat etc.) and organs (liver, brain, breast etc.) as well as other samples (Figure 1b-d). METASPACE includes data from all major ultra-high-resolution imaging MS technologies (MALDI-, DESI-, IR-MALDESI-, coupled with either Orbitrap and FTICR analyzers) and sample preparation protocols with various MALDI matrices represented (Figure 1e). Each dataset is provided with metadata (Supplementary Figure S1) describing sample, sample preparation, and mass spectrometry that enables searching, filtering or large-scale analysis. The knowledge base can be accessed either interactively through the webapp or programmatically (see “python-client” at https://github.com/metaspace2020/metaspace).

METASPACE knowledge base, as a large collection of data from various laboratories, helps understand the state of the art of imaging MS. A key advantage of imaging MS for untargeted analysis its high molecular coverage (or the number and types of molecules detectable in one experiment). Using METASPACE, we quantified the coverage as the number of annotations per dataset at FDR 10% and demonstrated that indeed, imaging MS has a high coverage detecting up to 10^3^ annotations with the most common numbers being in low hundreds (Figure 1f). We further quantified the dynamic range of imaging MS as the ratio between intensities of the most and least abundant annotated molecules and demonstrated that imaging MS has a dynamic range of up to 5 orders of magnitude with the most common values being around 10^3^ (Figure 1g). Another question, answering which otherwise requires extensive expert knowledge or literature search, is whether a particular molecule is detectable by imaging MS and how hard is it to detect it. Using METASPACE, one can query the molecule by either its name or molecular formula; the latter is recommended to avoid ambiguity in molecular namings. For example, the search for glutathione at FDR 10% returns 519 annotations from 18 laboratories (<u>link to glutathione images</u>). Interestingly, the histogram of the molecular prevalence shows that glutathione is one of the most prevalent molecules in METASPACE (Figure 1h). In comparison, the search for creatinine at FDR 10% returns 26 annotations only (<u>link to all creatinine images</u>) that indicates that both molecules are detectable but creatinine is less likely to be observed in the data. Further exploration of the metadata can show types of samples and technologies (e.g. source, polarity, or MALDI matrix) which contain or help detect a molecule of interest.

### Web app

The primary way to access METASPACE is through its web app for interactive exploration and sharing of spatial metabolomes (http://metaspace2020.eu/#/annotations). The web app can be opened in any web browser to submit data, access results, browse molecular images, overlay them with a microscopy image, perform molecular and metadata search, filter datasets by their metadata, organize own datasets into projects, as well as share projects, datasets or individual images using permalinks (Figure 1i-j). Not counting our team, the web app serves each month 10.000-25.000 pageviews or 600-1000 sessions with the most visitors being from the United States (30%), Germany (24%), France (7%), and the UK (7%) (Supplementary Figure S2). Figure 2 summarizes key scenarios of exploring metabolite images in METASPACE by illustrating them with images of various intestinal metabolites from whole-body animal tissue sections. Given a molecule of interest (e.g. taurocholic acid, a bile acid solubilizing fats in intestines; Figure 2a) one can browse other molecular images from the same dataset to search for co-localized ones (e.g. arginine, a known intestinal metabolite; Figure 2b). Next, one can visualize structurally-related molecules in the same dataset (e.g. arginyl-aspartate; Figure 2c).

**Figure 2.**
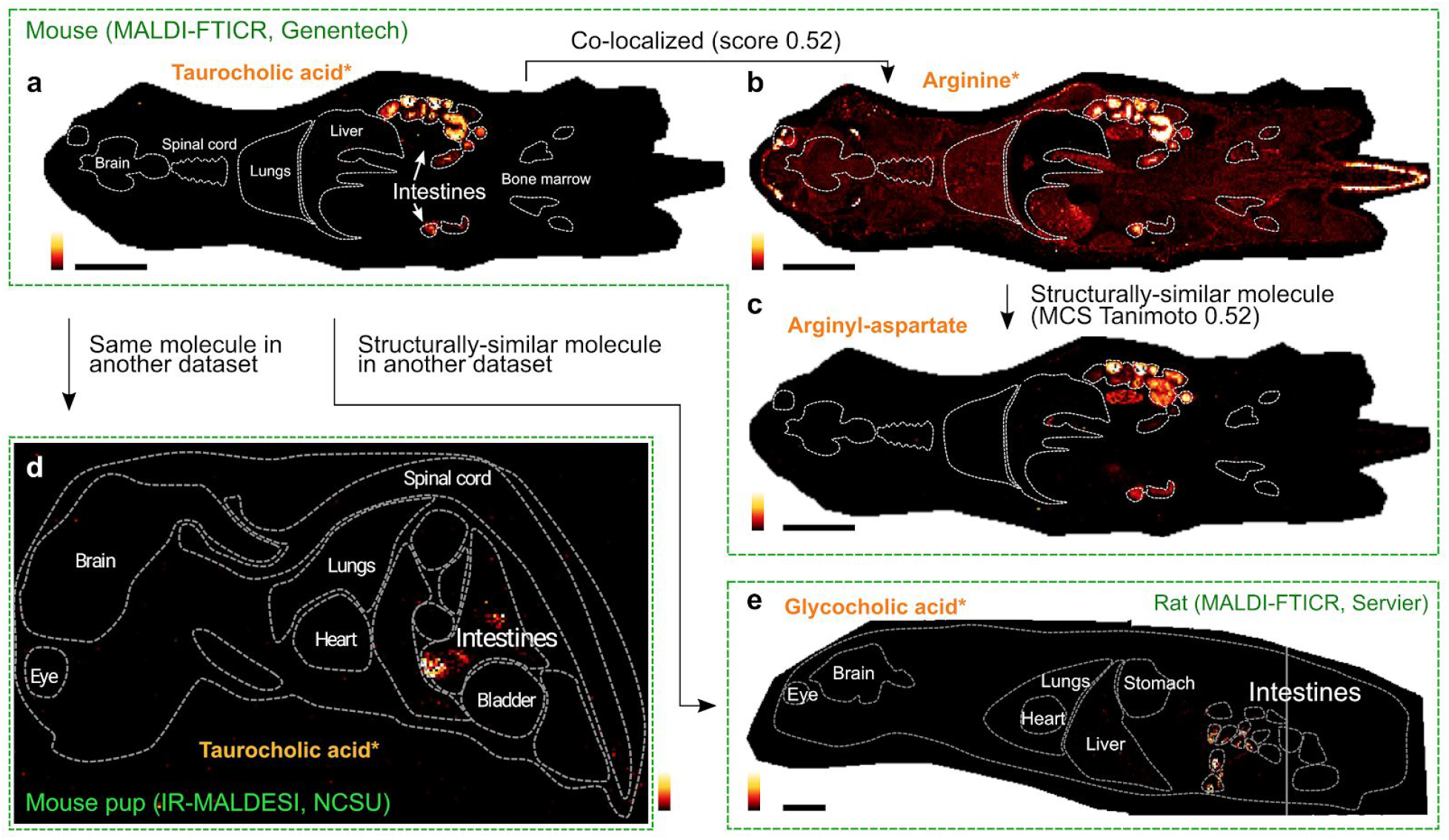
Summary of key scenarios of exploratory analysis of metabolite images in the METASPACE knowledge base as illustrated with various intestinal metabolites in whole-body animal model sections. **A.** Taurocholic acid, a bile acid solubilizing fats in intestines, exhibit respective localization. **B.** Arginine, a known intestinal metabolite, can be found in the same dataset by its co-localization with taurocholic acid. **C.** Arginyl-aspartate dipeptide, a molecule structurally similar to arginine, shows similar localization thus potentially indicating arginine metabolism in intestines. **D.** Intestinal localization of tarocholic acid in another dataset provides an additional confirmation of its association with this anatomic region. **E.** Glycocholic acid, another bile acid, also shows localization in intestines of a rat. Data contributed by Sheerin Shahidi-Latham, Genentech (<u>link</u>), Mark Bokhart, North Carolina State University (<u>link</u>), and Mathieu Gaudin, Servier (<u>link</u>). Co-localization score as described in Methods. MCS Tanimoto stands for the Maximum Common Subgraph Tanimoto distance between the molecular structures.

To confirm an association between a molecule of interest (e.g. taurocholic acid) and an anatomical or histological region, one can visualize its localization in another dataset (e.g. intestinal localization of taurocholic acid in another dataset; Figure 2d). Further exploration of other datasets can reveal molecules which are structurally-similar yet associated with the same anatomical region (e.g. glycocholic acid, a bile acid, shows intestinal localization in a rat whole-body tissue section; Figure 2e).

### Exploring spatial metabolomes on the levels of tissues and organs

In a case study, we explored how imaging content of METASPACE can generate or support hypotheses about localization of molecules within particular organs. We started the analysis with investigating data from whole-body animal model tissue sections originally used for drug development. For validation, we prepared, imaged and examined an organs-mimicking tissue section containing six wells embedded in gelatine, each contained homogentated mouse tissue from one of six organs (liver, heart, muscle, lung, kidney, and brain) following an earlier published protocol ^23^.

First, we investigated molecular images of glutathione (GSH), a major endogenous antioxidant with a high prevalence in METASPACE (Figure 3a). In mouse and rat whole-body sections, GSH is localized predominantly in the liver as well as in the salivary gland, lung, intestinal wall, and testis. Interestingly, GSH exhibited a distinct localization in the eye in a rat whole-body section. Analysis of the organs-mimicking tissue section confirmed the high abundance in the liver as compared to the other organs with traces in the lung and heart. This corroborates reports that GSH is mainly synthesized in liver ^24^ with a high (up to 7 mM) concentration in this organ ^25^. GSH is known to be abundant in testis with roles ranging from antioxidative protection of germ cells to contributions to protein synthesis and meiosis ^26^. GSH in the eye supports hydration and protects against chemical and oxidative stress ^27^. Next, we investigated the spatial aspects of the balance between GSH and oxidized glutathione (GSSG) that is critical for homeostasis. In the mouse whole-body section, compared to GSH, GSSG has a stronger association with the liver. The organs-mimicking tissue shows GSSG in the liver only. When comparing localizations of GSH and GSSG in a rat liver section analyzed at a high spatial resolution (20 μm), GSH and GSSG exhibit strikingly different and complementary localizations (Figure 3e). These images likely discern localization of GSH within cells where its synthesis and oxidation take place, and extracellular localization of produced GSSG pumped to the extracellular compartment ^25, 28^.

**Figure 3.**
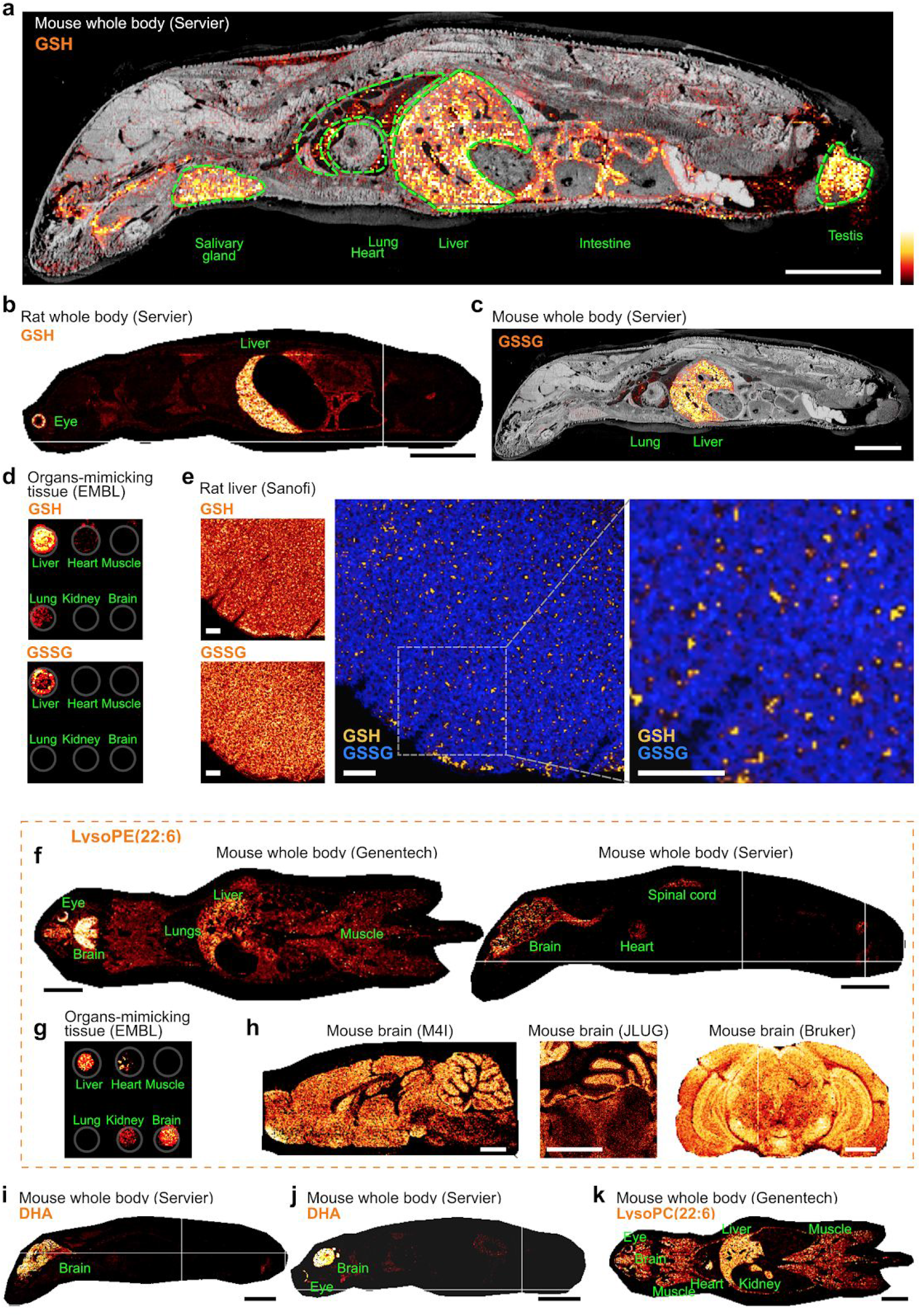
Two case studies exploring spatial metabolomes on the levels of tissues and organs for glutathione (**A-E**) and lysophosphatidylethanolamine LysoPE(22:6), an esterified form of docosahexaenoic acid (DHA). Glutathione (GSH) is localized in the liver, salivary gland, testis, intestines, and lung as shown in the mouse (**A**) and rat (**B**) whole-body sections and organs-mimicking tissue (**D**). **C.** Oxidized glutathione (GSSG) in localized predominantly in the liver. **E.** The data of higher spatial resolution for the rat liver shows different and complementary localizations of GSH and GSSG likely reflecting expected intracellular and extracellular concentrations of those; scale bar 500 μm. **F-H.** LysoPE(22:6) is localized in the brain, eye, spinal cord, liver, and heart of mouse whole-body sections (**F**) and organs-mimicking tissue (**G**). **H.** Images of higher spatial resolution show LysoPE(22:6) localization within the brain gray matter. **I-J.** Images for DHA high abundance in the brain and eye of mouse whole-body sections. **K.** LysoPC(22:6), another lipid storing DHA, is less specific to the brain. Data contributed by Mathieu Gaudin, Servier (<u>A</u>, <u>B</u>, <u>C</u>, <u>F</u>, <u>I</u>, <u>J</u>,), Cristina Gonzalez, EMBL (<u>D</u>, <u>G</u>), Marina Reuter, Sanofi (<u>E GSH</u>, <u>E GSSG</u>), Sheerin Shahidi-Latham, Genentech (<u>F</u>, <u>K</u>), Shane Ellis, M4I (<u>H</u>), Mario Kompauer, JLU Giessen (<u>H</u>), Michael Becker, Bruker (<u>H</u>). Scale bar for **a-c**, **f, i-k** 10 mm; scale bar for h 1 mm. Every images has its own intensity scale.

Next, we investigated molecular images of lysophosphatidylethanolamine LysoPE(22:6). This lipid, sometimes called LysoPE-DHA, is particularly interesting because it is an esterified docosahexaenoic acid (DHA), a predominant omega-3 polyunsaturated fatty acid with numerous functions in cognition, cancer, inflammation, and immunity ^29^. LysoPE(22:6) exhibit distinct localizations within the brain of the mouse whole-body sections, with traces in the eye, liver, heart, and spinal cord (Figure 3b). The organs-mimetic tissue also shows the highest concentration of LysoPE(22:6) in the brain with lower intensities in the liver, kidney, and heart. The images of a higher spatial resolution from different laboratories reproducibly associate LysoPE(22:6) within the grey matter. This is in line with the reports that DHA is the most abundant polyunsaturated fatty acid in the brain ^30^ and that upon crossing the blood-brain-barrier it is rapidly incorporated into phospholipids in particular into phosphatidylethanolamines ^31^. Molecular images of DHA in whole-body sections confirm this and also highlight its localization within retina, where DHA is known to play structural and protective roles ^32^. We also examined molecular images of lysophosphatidylcholine LysoPC(22:6) or LysoPC-DHA, another storage molecule of DHA (Figure 3b, also <u>more images in METASPACE</u>). Interestingly, the localization of LysoPC-DHA is drastically different from LysoPE-DHA with intensities not only throughout the brain and eye, but also in the muscles, liver, kidney, and heart. One could hypothesize that this is associated with the role of LysoPC-DHA as a carrier of DHA ^33^. However, LysoPC-DHA was not detected in the blood vessels (cf. the <u>carnitine localization</u> in the same section) that may indicate another prevalent function of LysoPC-DHA in these organs.

### Large-scale analysis of spatial metabolomes

Using the METASPACE molecular network, we demonstrate how large-scale analysis of big semi-structured data from METASPACE helps infer biologically relevant knowledge (Figure 4, Supplementary Figure S3). The organisation of the network in two distinct parts, each containing datasets collected using either negative or positive ion polarity (Figure 4a, Supplementary Figure S4), suggests that the ion polarity is a major driver of the molecular content in imaging MS. The Cohen’s kappa analysis confirmed the ion polarity to be indeed the most predictable factor with the next factors being the contributing laboratory and the organ (kappa of 0.95, 0.85 and 0.77, respectively; Supplementary Table S1). This is in line with other untargeted mass spectrometry analyses reporting that 90% of small molecules are detectable only in one ion polarity mode ^34^ Note that in METASPACE the metadata factors can be confounded and should be interpreted with caution (e.g. data for the lymph node is so far contributed by one laboratory only). However, for some organs, in particular for the brain, METASPACE already contains a sufficient number of samples allowing to differentiate it from other organs, as shown in a subnetwork of the brain samples from 26 laboratories (Figure 4b).

**Figure 4.**
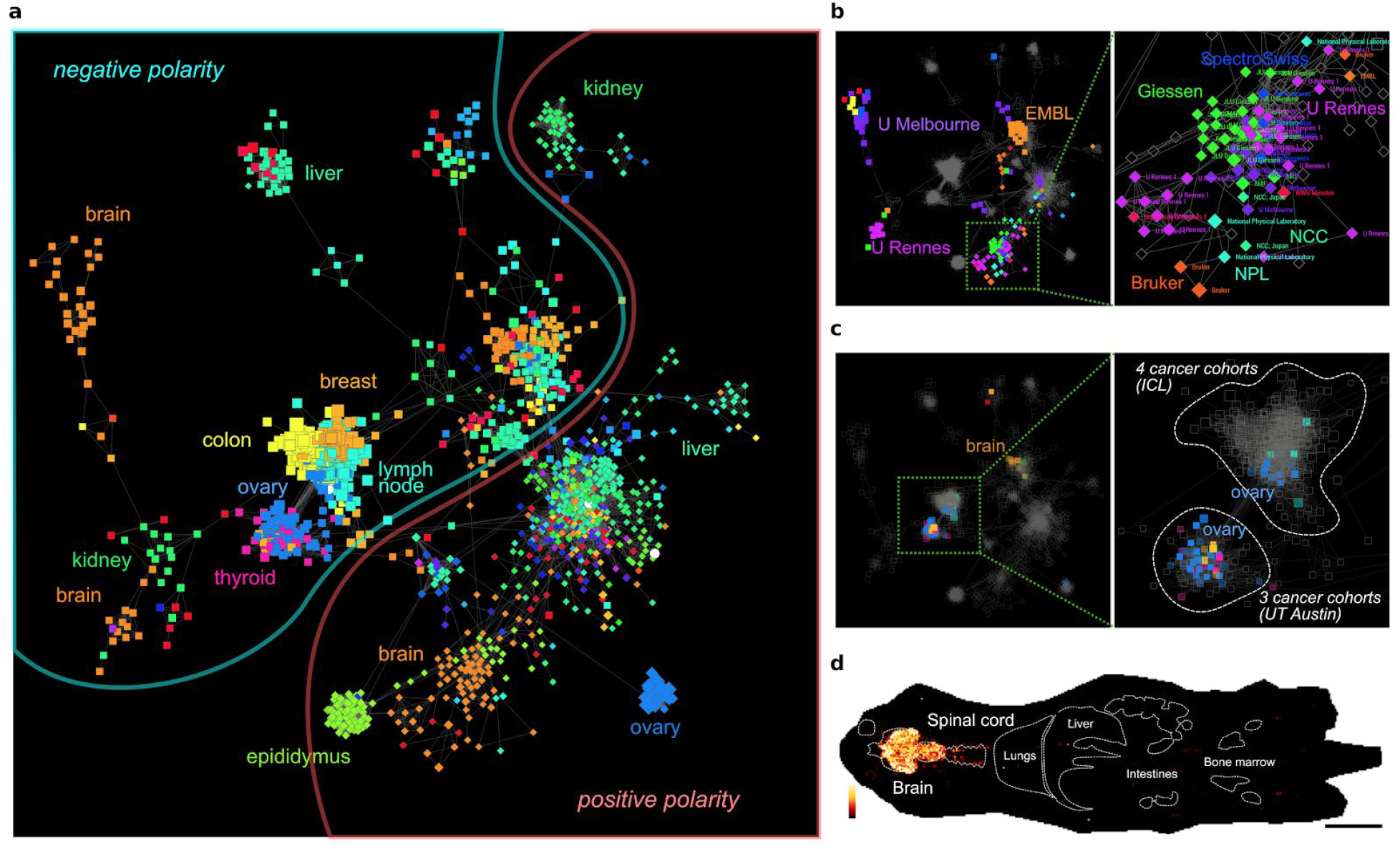
Large-scale analysis of spatial metabolomes in METASPACE. **A.** METASPACE molecular network showing all datasets organized by their molecular similarity. A node represents one dataset. An edge connects two datasets with the molecular similarity greater than 0.35. The datasets are colored by the sampled organ. **B.** Subnetwork associated with the brain representing data from 27 laboratories. **C.** Subnetwork of datasets containing N-acetylaspartic acid. **D.** Localization of N-acetylaspartic acid in a mouse whole-body section; scale bar 10 mm. See Supplementary Figures S3-S6 for larger views. Data in D contributed by Sheerin Shahidi-Latham, Genentech (<u>link</u>).

Using tf-idf modeling, we found molecules molecular markers characteristic and unique for the brain which were: 3-O-Sulfogalactosylceramide (d36:0), phosphatidylethanolamine PE(P-38:1), phosphoserine PS(36:1), gangliosides GM1 (38:1) and (36:0), and cholesteryl esters (Supplementary Table S2). We furthermore performed an enrichment analysis to identify characteristic molecular classes. For data analyzed in the negative ion mode it was sulfatides, phosphatidylinositols, LysoPEs, PEs, monoacylglycerophosphates (LysoPA and CPA), PSs, and phosphatidylglycerols among other classes (Supplementary Table S3). For the positive ion mode these were sphingomyelins and other phosphosphingolipids, and phosphatidylglycerophosphates (Supplementary Table S4). Interestingly, the same diacylglycerophosphates (phosphatidic acids PA(32:0), PA(36:4), PA(34:2), PA(34:1), PA(36:2)) were detected and found to be characteristic in both modes. Phosphatidylcholines were found characteristic in both modes with ca. two times higher enrichment score and number of detections in the positive mode. We also found organ-specific markers for kidney, lung, liver, and eye (see Supplementary Table S2). The markers for kidney were: L-carnitine (or an isomer), betaine (or isomers), 3-O-sulfogalactosylceramide (d34:1), cholesterol sulfate, uridine (or pseudouridine), and ganglioside GM2 (d44:0). The markers for liver were: linoleic acid (or conjugated linoleic acid), diacylglycerol DG(34:1), and a bile acid. With the continuing growth of METASPACE, we expect to obtain enough data to create comprehensive organ-specific molecular atlases for these and other organs.

The METASPACE molecular network can help explore associations between metabolites and disease. Figure 4c shows a subnetwork showing only datasets containing N-acetylaspartic acid (NAA). NAA is the second most abundant amino acid in the brain ^35^ that is illustrated with its localization in a whole-body mouse section (Figure 4d). Interestingly, NAA was detected not only in the brain, but also in two ovarian cancer cohorts: by the Takats lab, Imperial College London, and the Eberlin lab, University of Texas Austin. This, together with the fact that NAA was not detected in other cancer cohorts by those laboratories (Figure 4c), indicates its potential association with the ovarian cancer. This corroborates a recent report associating NAA with the reduced survival of patients with the high-grade serous ovarian cancer ^36^.

### Screening for drugs and their metabolites

METASPACE helps annotate and explore images not only of endogenous but also exogenous molecules, particularly drugs or other bioactive compounds as required in drug development. As of January 2019, 101 datasets were publicly shared by seven pharma companies (<u>link to all datasets</u>). As a case study, we explored a published dataset which was originally used for spatial quantitation of drugs in a rat liver ^14^(<u>dataset link</u>, Figure 5). Using METASPACE, we reproduced the original results ^14^ showing erlotinib in the liver of the cassette-dosed animals and olanzapine in both the cassette- and discrete-dosed animals. Moreover, when searching for metabolites of olanzapine from the DrugBank database ^18^ (Figure 5b), we discovered metabolites of olanzapine (isomeric 7-hydroxyolanzapine and 2-hydroxymethylolanzapine as well as 4’-N-desmethylolanzapine) which were not reported in the original publication (Figure 5c). Their presence in the liver sections but not in the mimetic tissue with drug standards confirms the expected *in vivo* metabolism of olanzapine. These images could be missed either due to targeted search for olanzapine only, or due to smaller intensities of the metabolites compared to olanzapine (by one and two orders of magnitude, respectively). This case study illustrates the potential of METASPACE in supporting drug development as well as creating additional value from the contributed public data.

**Figure 5.**
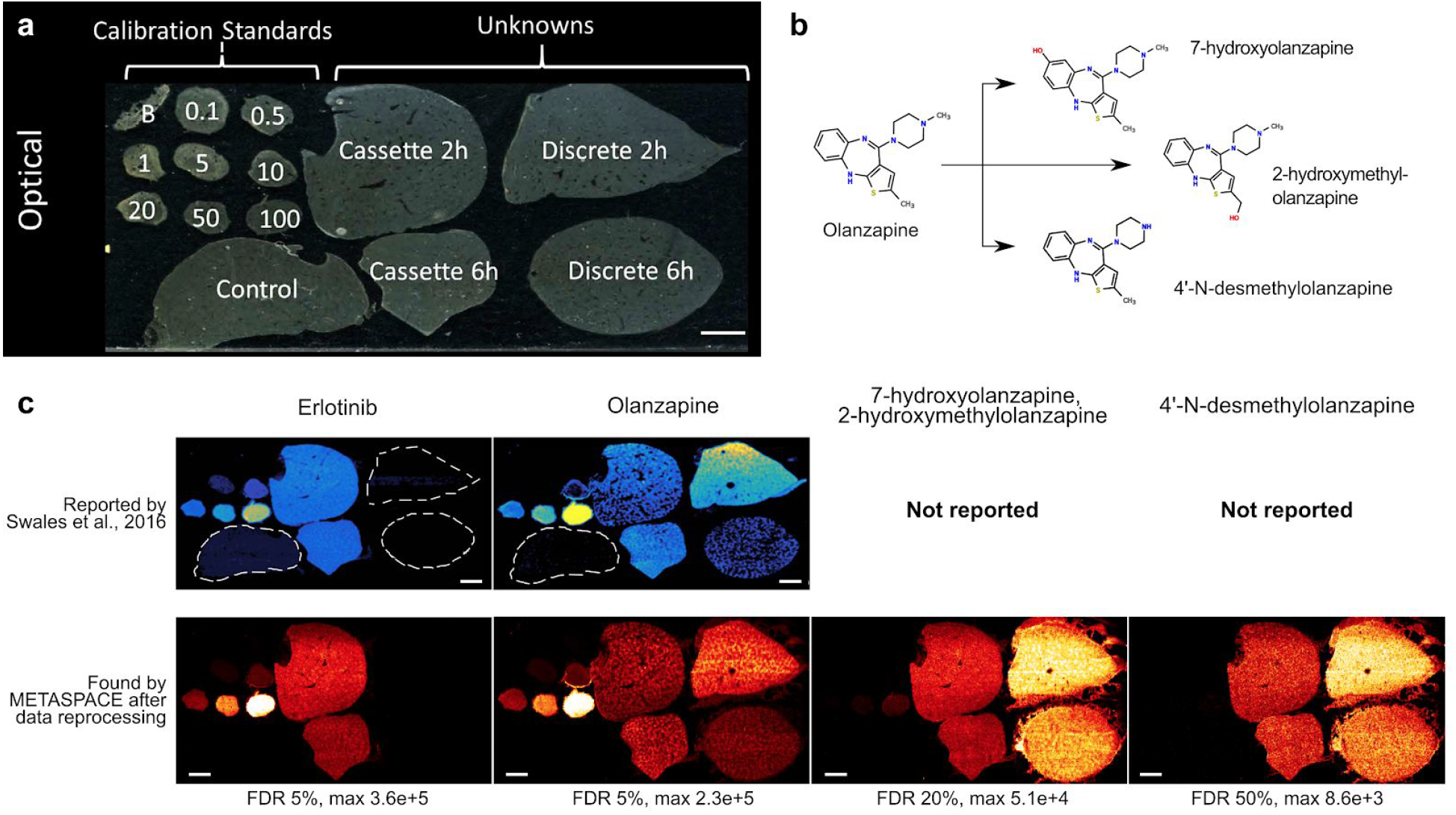
METASPACE helps localizing drug and drug metabolites in tissues. **A**: Optical image of liver sections from five animals including control, cassette dosage 2h-post and 6h-post, discrete (olanzapine only) 2h-post and 6h-post as well as a liver mimetic tissue section with drug standards; adapted from the original publication ^14^ **B**: Olanzapine metabolism according to DrugBank (https://www.drugbank.ca/drugs/DB00334). **C**: Molecular images reported in the original publication^14^ and those obtained by METASPACE show that METASPACE detected not only olanzapine but also its *in vivo* metabolites. Data contributed by John Swales and Nicole Strittmatter, AstraZeneca (<u>link</u> to olanzapine and its metabolites images). Scale bar 20 mm.

## Discussion

Here, we presented METASPACE, a community-populated knowledge base of spatial metabolomes already encompassing over 3200 public datasets. We illustrated how METASPACE facilitates metabolite annotation for imaging MS, reveals the state of the art of the imaging MS technology, enables large-scale analysis of spatial metabolomes, and supports drug development. METASPACE, although not yet published, was already used in several publications^37–43^ and highlighted in numerous reviews that indicates the significance and uniqueness of this resource for the imaging MS field. With the current rate of growth of 1000 datasets a year, its molecular and biological coverage will soon increase enough to provide spatial metabolomics atlases for various tissues, organs, and organisms. Our further steps include improvements of the engine to explore the unannotated “dark matter” of the already contributed datasets, implementation of various data analysis strategies in particular illustrated in this manuscript and supporting statistical and pathways analyses, further integration with repositories and resources for metabolomics ^44^ and other -omics, and engagement of computer scientists to optimize computations and build new services on top of METASPACE. This will multiply the positive effect of public data and support spatial metabolomics guide discoveries in health and disease.

## Online Methods

### Annotation engine

The annotation engine is a high-performance implementation of our computational method for metabolite annotation published earlier ^13^. The engine design follows the Resilient Distributed Datasets paradigm and is implemented using the Apache Spark framework and the pySpark binding. The engine is deployed onto the Amazon Web Services cloud and leverages a cluster of several cloud instances, usually three AWS c4.2xlarge instances each with 8 vCPU and 15 GiB memory. The engine is open-source, available on GitHub (https://github.com/metaspace2020/metaspace, repository “engine”) and developed by a team of professional software developers following modern software practices including Agile development, unit testing, integration testing, and code review. The engine takes as input a centroided imaging MS dataset in the imzML format ^17^, a molecular database including molecular formulas, and ion adducts. For each ion, the engine predicts its fine isotopic structure considering the resolving power provided upon submission and uses the 3ppm tolerance for pulling m/z-images of four principal peaks from the imzML file. Several molecular databases are available for the metabolomics and lipidomics annotation. The Human Metabolome Database, HMDB, ^18^ is used as the default database when screening for metabolites in human and animal models samples. We also prepared an HMDB-endogenous version of HMDB which contains only those molecules which are labelled in HMDB as endogenous (the HMDB property “Source” is equal to “Endogenous”) to reduce potential false positives when screening for endogenous metabolites. Another imported database, Chemical Entities of Biological Interest, ChEBI, ^19^ is recommended for integrating results with other -omics, since its chemical entities are organized in a chemical ontology linked to GeneOntology, UniProt and the BRENDA database of enzymes. For annotating lipids, we imported the LIPID MAPS ^20^ and SwissLipids ^21^ databases. Following users requests, we also imported the species-specific databases such as PAMDB for *Pseudomonas aeruginosa*^45^ metabolites and ECMDB for *Escherichia coli*^46^.

### Knowledge base and web application

The METASPACE knowledge base is a collection of public metabolite annotation results contributed by users. At the heart of the knowledge base is a cloud software which integrates the following services operating individually and communicating with each other via REST APIs by using a GraphQL interface: 1) the metabolite annotation engine described earlier, 2) a service containing and providing molecular databases, 3) a web application for submitting data as well as browsing and sharing the results, and 4) API for programmatic access of the annotations and metadata (“engine”, “mol-db”, “webapp”, “python-client” repositories at https://github.com/metaspace2020/metaspace, respectively). An essential component of the knowledge base that ensures future re-use of data is the metadata contributed for each dataset during the submission (Supplementary Figure METADATA). The required metadata includes information about the sample (organism, organ, condition), sample preparation (sample stabilization, tissue modification, MALDI matrix, matrix spraying), mass spectrometry (source, analyzer, mass resolving power), and contributor contact information (laboratory, name, principal investigator). The storage of the metabolite annotation results is organized for sustainable handling of big data and includes a PostgreSQL relational database for consistent storage and an ElasticSearch search engine for fast querying and access. ElasticSearch was estimated to be at least 10x faster than PostgreSQL for our type and size of data. This high-performance access is especially beneficial, since the knowledge base encompasses a large number of annotations (over 6 million as of January 2019). For data upload, we use a big data upload service which restarts in case of an interrupted connection. The imzML files are stored on the Amazon S3 storage. The web app is implemented using the Vue.js JavaScript framework and enables data submission, annotations browsing, searching, or filtering, as well as visualization of ion images and other diagnostics information. The web app uses TypeScript to increase stability and improve our productivity when working on the codebase. The GraphQL, a data query and manipulation language, is used as the interface between the services, to provide an API for the web application, as well as for programmatic access to the knowledge base. RabbitMQ is used for queuing and synchronization of the services.

We provide data management capabilities, where a user needs to be authenticated for submission and handling their private data. Users from the same laboratory can be organized into groups by their group admins and share their private data with each other. Users can also organize their data in projects which can be both private and public. This allows users to organize data by associated experiments as well as to share datasets for publication by creating a project and sharing all the datasets from this project using a URL.

Regular snapshots of the knowledge base including all public annotation results are provided at https://github.com/metaspace2020/pixel-annot-export.

### Tf-idf modeling of datasets

For data mining, we represented METASPACE datasets as vectors in a molecular space. Inspired by natural language processing approaches to clustering, classification, and searching text documents^47^, we modelled METASPACE datasets as “documents” and molecular formulas annotated within the dataset as “words”. For each molecular formula annotated in a dataset, we calculated the *term frequency-inverse document frequency* (tf-idf) statistic which quantifies how characteristic the molecular formula is for this dataset compared to all other METASPACE datasets. The tf-idf statistic is defined as *tfidf*(*m, d, D*) = *tf*(*m, d*) * *idf*(*m, D*), where *m* is a molecular formula, d is a dataset, and *D* is the set of all METASPACE datasets. The “frequency” of each molecular formula *m* (“word” or “term”) in a dataset *d* (“document”) was defined to be inversely proportional to the FDR value *F* associated with *m* in d: *tf*(*m, d*) = 100 − *F*, where *F* takes values 5%, 10%, 20%, or 50%. We defined *idf*(*m, D*) as *log_n_* of the ratio between the number of all datasets *D* in METASPACE and the number of those datasets where the molecular formula *m* was annotated with FDR 50%. As a result, we represented each dataset as a tf-idf vector in the space of all molecular formulas annotated in METASPACE. The Python library gensim v2.3.0 was used for calculating the tf-idf vectors.

### Quantifying the influence of metadata parameters on the molecular content

We developed the following approach to quantify how much a metadata parameter (e.g. ion polarity, laboratory, organism, organ, or MALDI matrix) influences the molecular content of the associated data. First, for each metadata parameter *M* (e.g. organ), we considered its values (e.g. “brain”, “kidney”, etc). For each dataset, we found its first-nearest-neighbor (NN) dataset among all datasets according to the cosine similarity in the tf-idf space. This search resulted in the so-called NN-pairs of datasets. We calculated *p_NN_*, the percentage of the NN-pairs of datasets having the same value of the metadata parameter *M* (e.g. for a “brain” dataset its NN dataset is also a “brain” dataset). Next, for each dataset we randomly selected another dataset that resulted in the so-called random pairs. We calculated *p_random_*, the percentage of those random pairs whose values of the metadata parameter *M* coincide. As a measure of predictability of the metadata parameter *M* in all datasets, we used the Cohen’s kappa *x* = 1 − (1 − *p_NN_*)/(1 − *p_random_*) ^48^. The Cohen’s kappa is a enrichment-factor-like statistic which compares the observed accuracy vs the expected accuracy. The Cohen’s kappa compensates for a random chance and accounts for different numbers of parameter values and uneven group sizes thus producing values comparable between different metadata parameters. The Cohen’s kappa takes values from −1 to 1 and the higher is the value for a metadata parameter *M*, the higher is the predictability of this parameter from the data, or the higher is influence of the parameter *M* on the molecular content of the data.

### Data similarity network

We developed a molecular networking approach to represent all public METASPACE datasets as a network based on their molecular content. After the tf-idf modeling of all datasets, the molecular similarity between two datasets was defined as the cosine similarity between their tf-idf vectors. The datasets with the similarity greater than 0.35 were connected with edges and visualized in the Cytoscape software.

### Characteristic molecules

In order to find characteristic molecular formulas for a particular dataset, we considered its molecular formulas with the highest tf-idf values. To find characteristic molecular formulas for a group of datasets (e.g. for a particular organ), we considered formulas with the highest average values across the group tf-idf values.

### Molecular class enrichment analysis

In order to find molecular classes enriched in molecular formulas annotated in a group of datasets (e.g. those characteristic for a particular organ), we developed the following enrichment analysis based on the tf-idf modeling. We considered the ClassyFire molecular taxonomy which is available for all molecules in HMDB^49^. For each molecular class *C* and a group of datasets *G*, we defined the enrichment score as 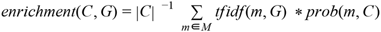, where |*C*| is the number of molecular formulas in class *C*, *M* is the set of all molecular formulas in METASPACE, *tfidf(m, G)* is an average tf-idf score of molecular formula *m* over datasets in *G*, and *prob*(*m, C*) = |*si*(*m, C*)| * |*si*(*m*)|^−1^ is a probability that molecule *m* belongs to class *C* when accounting for its structural isomers, with |*si*(*m, C*)| being the number of structural isomers of molecule *m* in class *C* and |*si*(*m*)| being the number of structural isomers of molecule *m*.

### Co-localization score

In order to find ions co-localized within a dataset for an ion of interest, we used the following method. For ions annotated in the dataset with an FDR <= 50%, we applied a hotspot removal by quantile thresholding with the quantile 0.99. Then, we calculated a co-localization score as a pairwise cosine similarity between pixel-wise intensity vectors for each two ions in the dataset using the Python library *gensim* v2.3.0 and considered top ions most similar to the ion of interest.

