## Supplementary Figures for "METASPACE: A community-populated knowledge base of spatial metabolomes in health and disease"

### Supplementary Figures for the manuscript "METASPACE: A community-populated knowledge base of spatial metabolomes in health and disease"

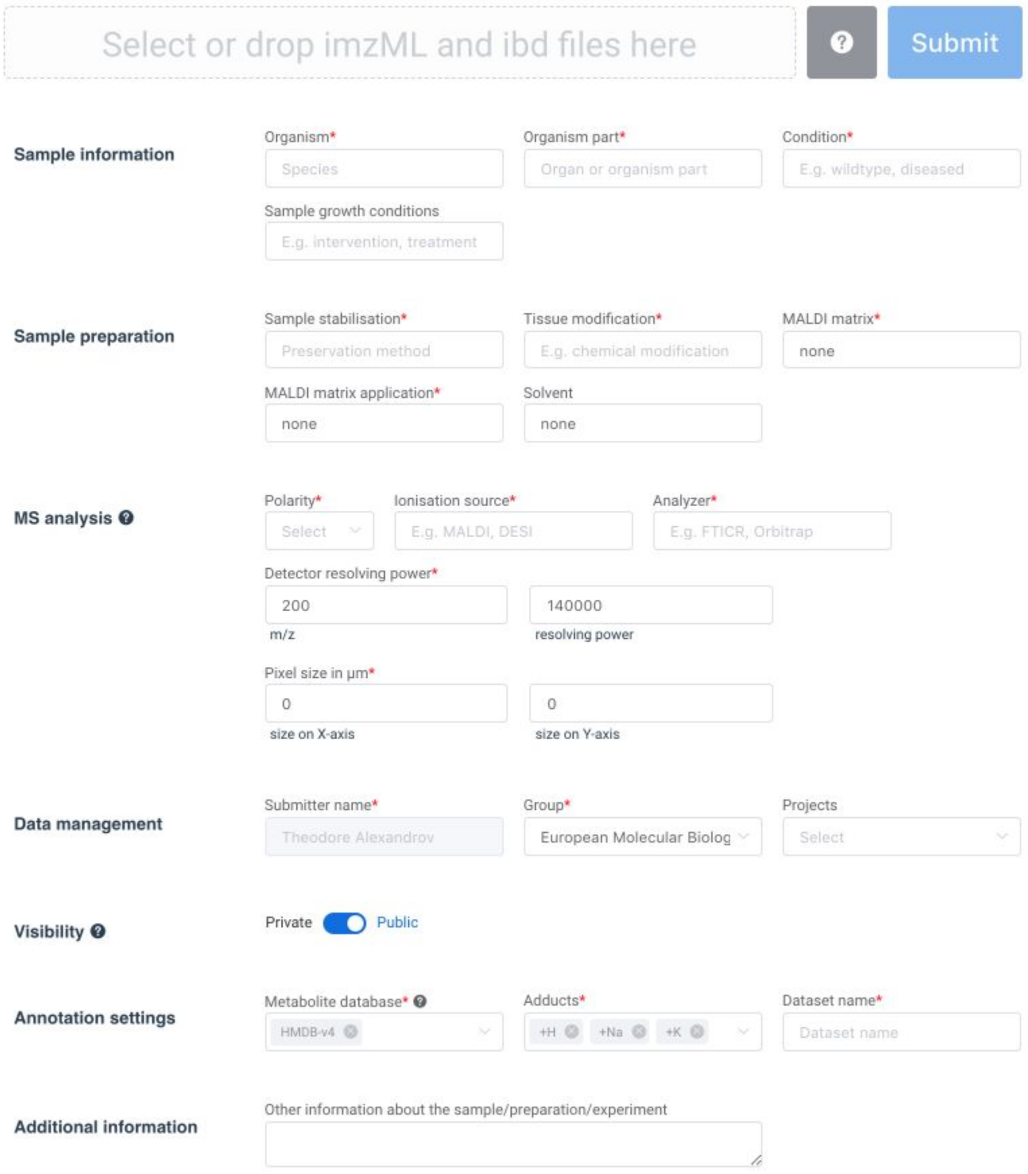


**Supplementary Figure S1.** A screenshot of the Upload page of the METASPACE web app that shows the metadata collected during submission. The required metadata is highlighted with a red star. For more information on the metadata, see "metadata" repository at https://github.com/metaspace2020/metaspace.

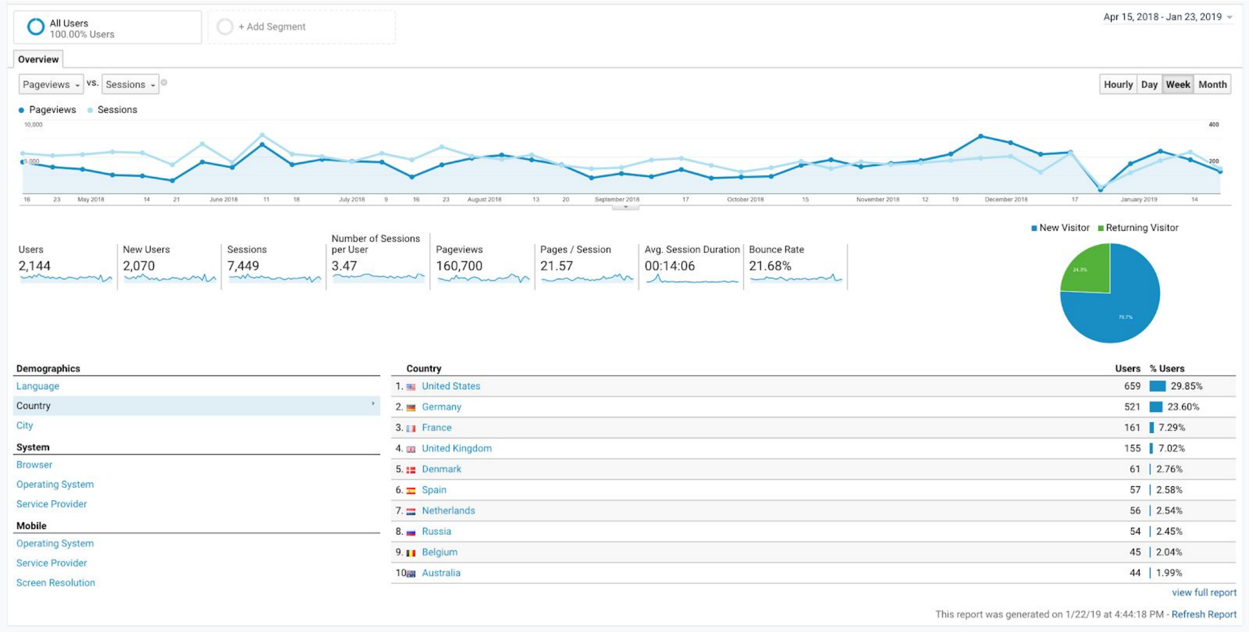


**Supplementary Figure S2.** A screenshot from the Google Analytics for the METASPACE webapp (http://metaspace2020.eu) showing the numbers of pageviews and sessions per month in 2018 as well as breaking down the visits by the countries.

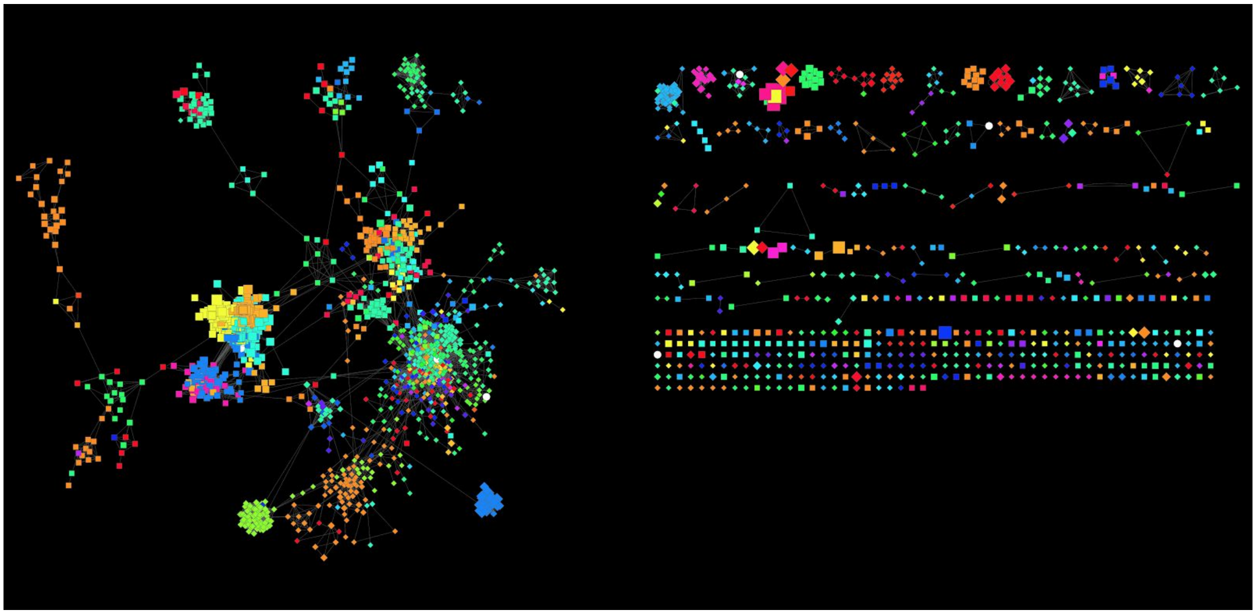

**Supplementary Figure S3.** The METASPACE network showing all datasets colored by the organ the tissue was sampled from. The connected part of the network on the left represents the majority of datasets and was shown in Figure 4a.

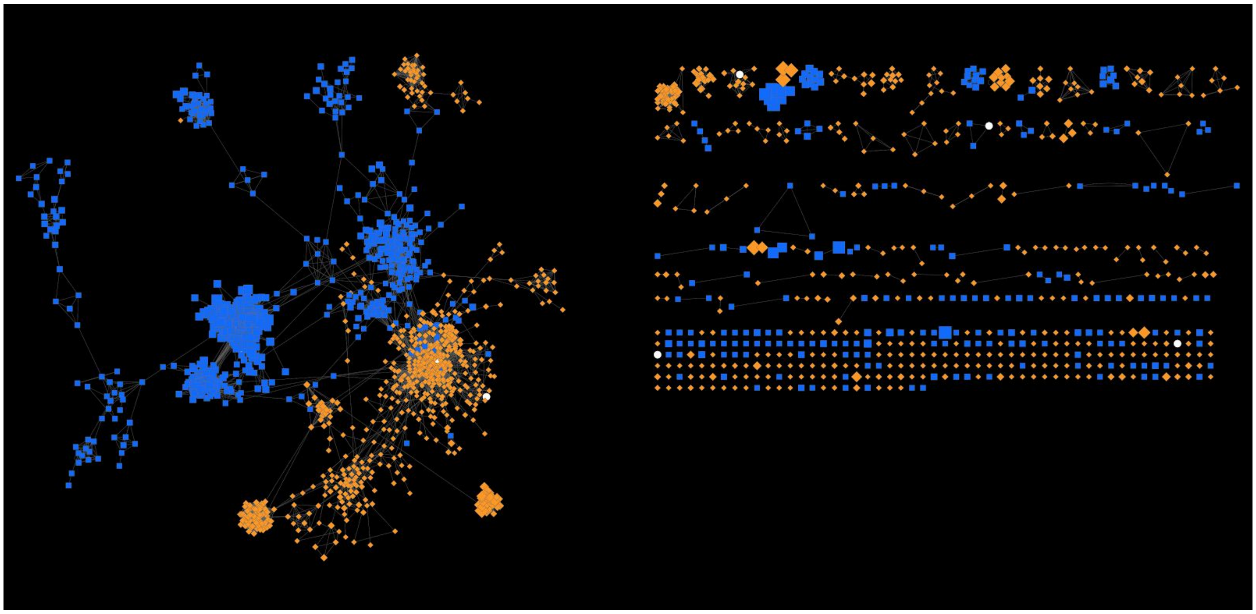

**Supplementary Figure S4.** The METASPACE network showing all datasets colored by the ion polarity mode used for the acquisition (blue and yellow for the negative and positive polarities, respectively). The connected part of the network on the left represents the majority of datasets and was shown in Figure 4a.

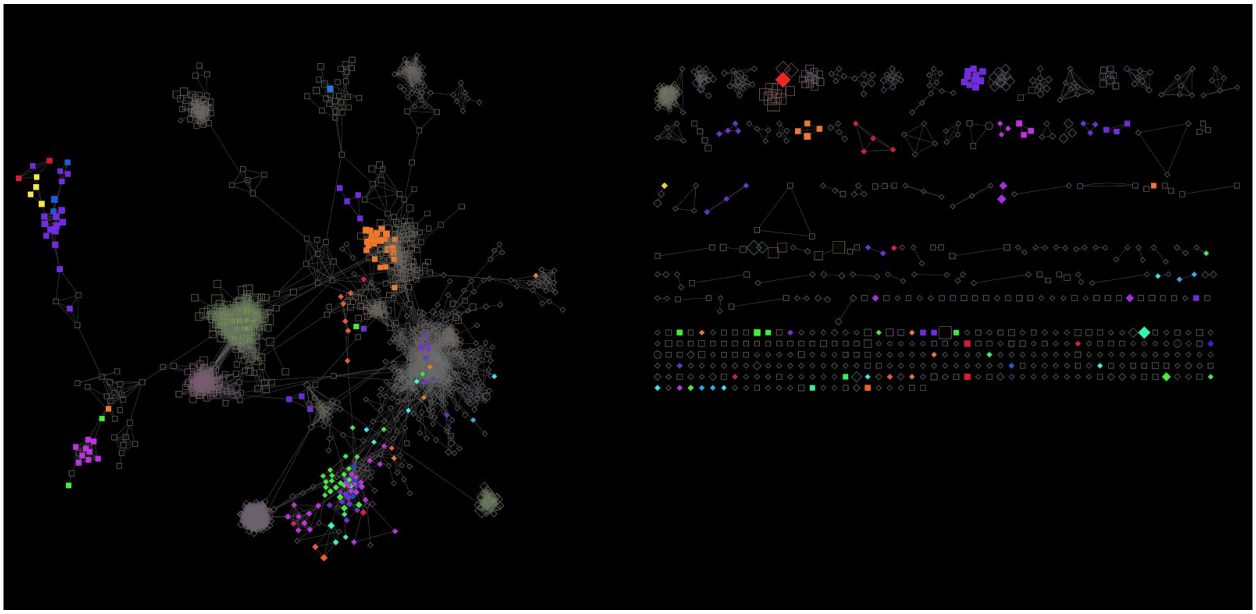

**Supplementary Figure S5.** The METASPACE network highlighting datasets from the brain and colored by the contributing laboratory. The connected part of the network on the left represents the majority of datasets and was shown in Figure 4b.

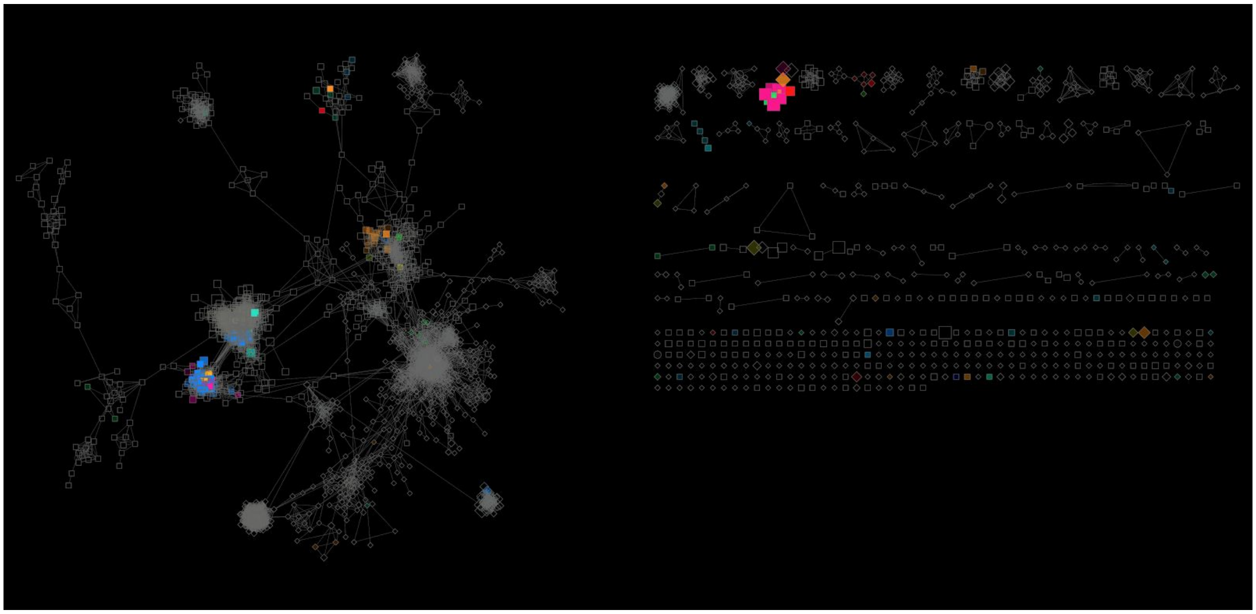

**Supplementary Figure S6.** The METASPACE network highlighting datasets where N-acetylaspartic acid was annotated and colored by the organ the tissue was sampled from. The connected part of the network on the left represents the majority of datasets and was shown in Figure 4c.
