## Supplementary Tables for "METASPACE: A community-populated knowledge base of spatial metabolomes in health and disease"

### Supplementary Tables for the manuscript "METASPACE: A community-populated knowledge base of spatial metabolomes in health and disease"

**Supplementary Table S1.** The values of Cohen's kappa for the metadata factors for all datasets in METASPACE represent predictability of these factors from the molecular content of the data.

| Factor | Td-idf similarity | Random similarity | Cohen's kappa |
|---|---|---|---|
| Ion polarity | 0.9730 | 0.4974 | 0.9463 |
| Lab | 0.8663 | 0.1055 | 0.8506 |
| Organ | 0.7699 | 0.0571 | 0.7560 |
| Organism | 0.8013 | 0.2785 | 0.7246 |
| MALDI matrix | 0.7406 | 0.2075 | 0.6727 |

**Supplementary Table S2.** See the file "*Supplementary Table S2. Organ-specific markers.pdf*"

**Supplementary Table S3.** See the file "*Supplementary Table S3. Molecular class enrichment analysis, brain, negative mode.xlsx*"

**Supplementary Table S4.** See the file "*Supplementary Table S4. Molecular class enrichment analysis, brain, positive mode.xlsx*"
