## Supplementary Table S2 for "METASPACE: A community-populated knowledge base of spatial metabolomes in health and disease"

**Supplementary Table. Organ-specific markers for brain, kidney, lung, liver, and eye, discovered by mining the METSPACE knowledge base.** We calculated the markers by considering all datasets in METASPACE. For each molecule, we calculated its average tf-idf value for all datasets sampled from this organ ("Tf-idf"). We show only the average tf-idf values above 0.05. As an additional measure of reproducibility of each marker in each organ, we show by how many labs it was detected in data from the organ with an FDR <= 10% ("#Labs"). In order to reproduce the findings as well as show the relative intensities of the markers within the same molecular ion image, when detected we show images for a mimetic tissue model of homogenated organs from mouse. Each of the six wells represents one of the homogenated organs: liver, heart, muscle, lung, kidney, and brain.

| Molecular formula | Molecule | Brain | | Kidney | | Lung | | Liver | | Eye | | Mimetic organs image |
|---|---|---|---|---|---|---|---|---|---|---|---|---|
| | | Tf-idf | # Labs | Tf-idf | # Labs | Tf-idf | # Labs | Tf-idf | # Labs | Tf-idf | # Labs | |
| C45H78NO8P | PE(40:6) | 0.17 | 16 | 0.06 | 5 | | | | | 0.26 | 1 | 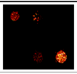 link |
| C42H81NO11S | 3-O-Sulfogalactosylceramide (d18:1/18:0) | 0.15 | 9 | | | | | | | | | 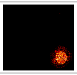 link |
| C43H82NO7P | PE(P-38:1) | 0.12 | 12 | | | | | | | | | 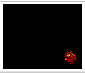 link |
| C40H80NO8P | PC(32:0) | 0.11 | 19 | 0.07 | 7 | 0.10 | 1 | 0.06 | 6 | 0.18 | 1 | 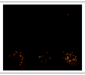 link |
| C39H73O8P | PA(36:2) | 0.10 | 18 | | | | | 0.09 | 6 | | | 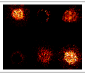 link |
| C27H44NO7P | LysoPE(22:6) | 0.09 | 14 | | | 0.05 | 1 | | | | | 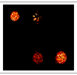 link |
| C42H80NO10P | PS(36:1) | 0.09 | 13 | | | | | | | | | 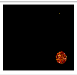 link |
| C75H135N3O31 | Ganglioside GM1 (18:1/20:0) | 0.09 | 2 | | | | | | | | | |
| C73H131N3O31 | Ganglioside GM1 (18:1/18:0) | 0.08 | 2 | | | | | | | | | |
| C39H79N2O6P | SM(d18:0/16:1) / Palmitoyl sphingomyelin | 0.06 | 12 | 0.05 | 7 | 0.08 | 1 | 0.06 | 6 | 0.06 | 1 | |
| C8H20NO6P | Glycerophosphocholine | 0.06 | 9 | 0.18 | 6 | 0.07 | 1 | | | | | |
| C43H70O3 | Cholesteryl ester | 0.05 | 5 | | | | | | | | | |
| C7H15NO3 | L-Carnitine, Malonyl-Carnitin | | | 0.13 | 3 | | | | | | | |
| C9H17NO4 | L-Acetylcarnitine / N-lactoyl-Leucine | | | 0.12 | 4 | 0.16 | 1 | | | | | |
| C5H11NO2 | Betaine / L-Valine / N-Methyl-a-aminoisobutyric acid / 5-Aminopentanoic acid | | | 0.11 | 4 | | | | | | | |
| C40H77NO11S | 3-O-Sulfogalactosylceramide (d18:1/16:0) | | | 0.11 | 5 | | | | | | | 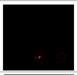 link |
| C27H46O4S | Cholesterol sulfate | | | 0.09 | 5 | | | | | | | 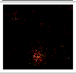 link |
| C29H47NO4 | Acylcarnitine (Docosa-4,7,10,13,16-pentaenoyl carnitine / Clupanodonyl carnitine) | | | 0.06 | 5 | 0.11 | 1 | 0.05 | 4 | 0.09 | 1 | |
| C10H12N4O5 | Inosine / Allopurinol riboside | | | 0.06 | 6 | | | | | | | 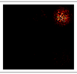 link |
| C9H12N2O6 | Uridine / Pseudouridine | | | 0.06 | 4 | | | | | | | |
| C76H140N2O26 | Ganglioside GM2 (d18:0/26:0) | | | 0.06 | 2 | | | | | | | |
| C40H78NO8P | PC(32:1) | | | | | 0.07 | 1 | 0.05 | 5 | | | |
| C10H26N4 | Spermine | | | | | 0.06 | 1 | | | | | |
| C10H13N5O4 | Adenosine / Deoxyguanosine | | | | | 0.05 | 1 | | | | | |
| C18H32O2 | Linoleic acid / conjugated linoleic acid | | | | | | | 0.07 | 6 | | | 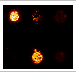 link |
| C37H70O5 | DG(34:1) | | | | | | | 0.07 | 3 | | | |
| C26H45NO6S | Tauroursodeoxycholic acid / Taurodeoxycholic acid / Taurochenodesoxycholic acid | | | | | | | 0.05 | 4 | | | 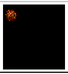 link |

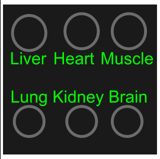
